# Acoustic Remote Control of Bacterial Immunotherapy

**DOI:** 10.1101/2021.03.25.434639

**Authors:** Mohamad H. Abedi, Michael S. Yao, David R. Mittelstein, Avinoam Bar-Zion, Margaret Swift, Audrey Lee-Gosselin, Mikhail G. Shapiro

**Affiliations:** Division of Biology and Biological Engineering, California Institute of Technology, Pasadena, CA, USA; Division of Engineering and Applied Sciences, California Institute of Technology, Pasadena, CA, USA; Division of Chemistry and Chemical Engineering, California Institute of Technology, Pasadena, CA, USA

## Abstract

Rapid advances in synthetic biology are driving the development of genetically engineered microbes as therapeutic agents for a multitude of human diseases, including cancer. In particular, the immunosuppressive microenvironment of solid tumors creates a favorable niche for systemically administered bacteria to engraft in the tumor and release therapeutic payloads. However, such payloads can be harmful if released in healthy tissues where the bacteria also engraft in smaller numbers. To address this limitation, we engineer therapeutic bacteria to be controlled by focused ultrasound, a form of energy that can be applied noninvasively to specific anatomical sites such as solid tumors. This control is provided by a temperature-actuated genetic state switch that produces lasting therapeutic output in response to briefly applied focused ultrasound hyperthermia. Using a combination of rational design and high-throughput screening we optimized the switching circuits of engineered cells and connected their activity to the release of immune checkpoint inhibitors. In a clinically relevant cancer model, ultrasound-activated therapeutic microbes successfully turned on *in situ* and induced a marked suppression of tumor growth. This technology provides a critical tool for the spatiotemporal targeting of potent bacterial therapeutics in a variety of biological and clinical scenarios.

## INTRODUCTION

Cell therapies are rapidly emerging as an exciting and effective class of technologies for cancer treatment^1–3^. Among the cell types being investigated for therapy, immune cells have excelled in the treatment of hematologic malignancies. However, their use in solid tumors has been hampered by their reduced ability to penetrate and function in the tumor’s immunosuppressive environment, especially within immune-privileged hypoxic cores^4–6^. Conversely, the reduced immune activity of some tumor cores creates a favorable microenvironment for the growth of certain bacteria, which can reach the tumors after systemic administration^7–9^. Capitalizing on their tumor-infiltrating properties, such bacteria can be engineered to function as effective cellular therapies by secreting therapeutic payloads to directly kill tumor cells or remodel the microenvironment to stimulate antitumor immunity^10–15^. However, the benefits of microbial therapy are often counterbalanced by safety concerns accompanying the systemic injection of microbes into patients with limited control over their biodistribution or activity^1,16,17^. This is especially important given the well-documented engraftment of circulating bacteria into healthy tissues such as the liver, spleen, and certain hypoxic stem cell niches^18–21^. To avoid damaging healthy organs, it is crucial that the therapeutic activity of microbes be targeted to tumors.

Among the available mechanisms to regulate microbial function, systemically administered chemical inducers^22,23^ are incapable of targeting a particular anatomical site. Meanwhile, light-induced control elements provide high spatiotemporal precision^24–26^, but are constrained by the poor penetration of light into intact tissues^27^. In contrast, temperature-based control elements provide a combination of spatiotemporal control and depth, since temperature can be elevated precisely in deep tissues using noninvasive methods such as focused ultrasound (FUS)^28–30^.

Indeed, it was recently demonstrated that FUS can be used in conjunction with temperature-dependent transcription factors to control the expression of bacterial genes^31^. However, these transcription factors operated in therapeutically irrelevant cloning strains of bacteria, had non-therapeutic outputs, and produced only transient activation unsuitable for tumor treatment, which typically requires weeks of therapeutic activity^10,11,22^.

Here we describe the development of FUS-activated therapeutic bacteria in which a brief thermal stimulus activates sustained release of anti-cancer immunotherapy. We engineer these cellular agents by adapting temperature-sensitive transcription factors to the tumor-homing probiotic species *E. coli* Nissle 1917 and designing gene circuits in which they control an integrase-based state switch^31,32^ resulting in long-term therapy production. To improve the safety and efficacy of these cells, we screen random and rationally designed libraries of gene circuit variants for constructs with minimal baseline activity and maximal induction upon thermal stimulation. We use the optimized gene circuits to express immune checkpoint inhibitors targeting CTLA-4 and PD-L1. In a mouse cancer model, we show that the resulting engineered microbes are reliably and chronically activated by a brief, noninvasive FUS treatment after systemic administration to release therapy and successfully suppress tumor growth.

## RESULTS

### Characterizing thermally responsive repressors in a therapeutically relevant microbe

To develop a temperature-actuated therapeutic circuit, we started with high-performance temperature-dependent transcriptional repressors, which actuate transient gene expression in response to small changes in temperature around 37 °C^31^. Since genetic elements tend to behave differently across cell types due to variations in protein expression and other aspects of the intracellular environment^33^, we first characterized the performance of these repressors in our chosen therapeutic chassis: *E. coli* Nissle 1917 (EcN). This bacterial strain is approved for human probiotic use and is commonly employed in microbial tumor therapies^10,34^. We selected three repressor candidates—TlpA39, wild-type TcI, and TcI42—as our starting points due to their desirable activation temperature thresholds of 39 °C, 38 °C, and 42 °C, respectively

To evaluate the performance of these candidates we designed reporter constructs where they regulate the expression of a green fluorescent protein (GFP) (**Fig. 1a**), transformed them into EcN cells, and measured the corresponding cell density-normalized fluorescence intensity as a function of temperature between 33 °C and 42 °C (**Fig. 1b**). As we were interested in using these bioswitches *in vivo*, we focused on the fold-change between the mammalian physiological temperature (37 °C) and an elevated temperature that can be used to trigger activation *in vivo* while minimizing thermal damage to local tissues (42 °C) (**Fig. 1c**).

**Figure 1.**
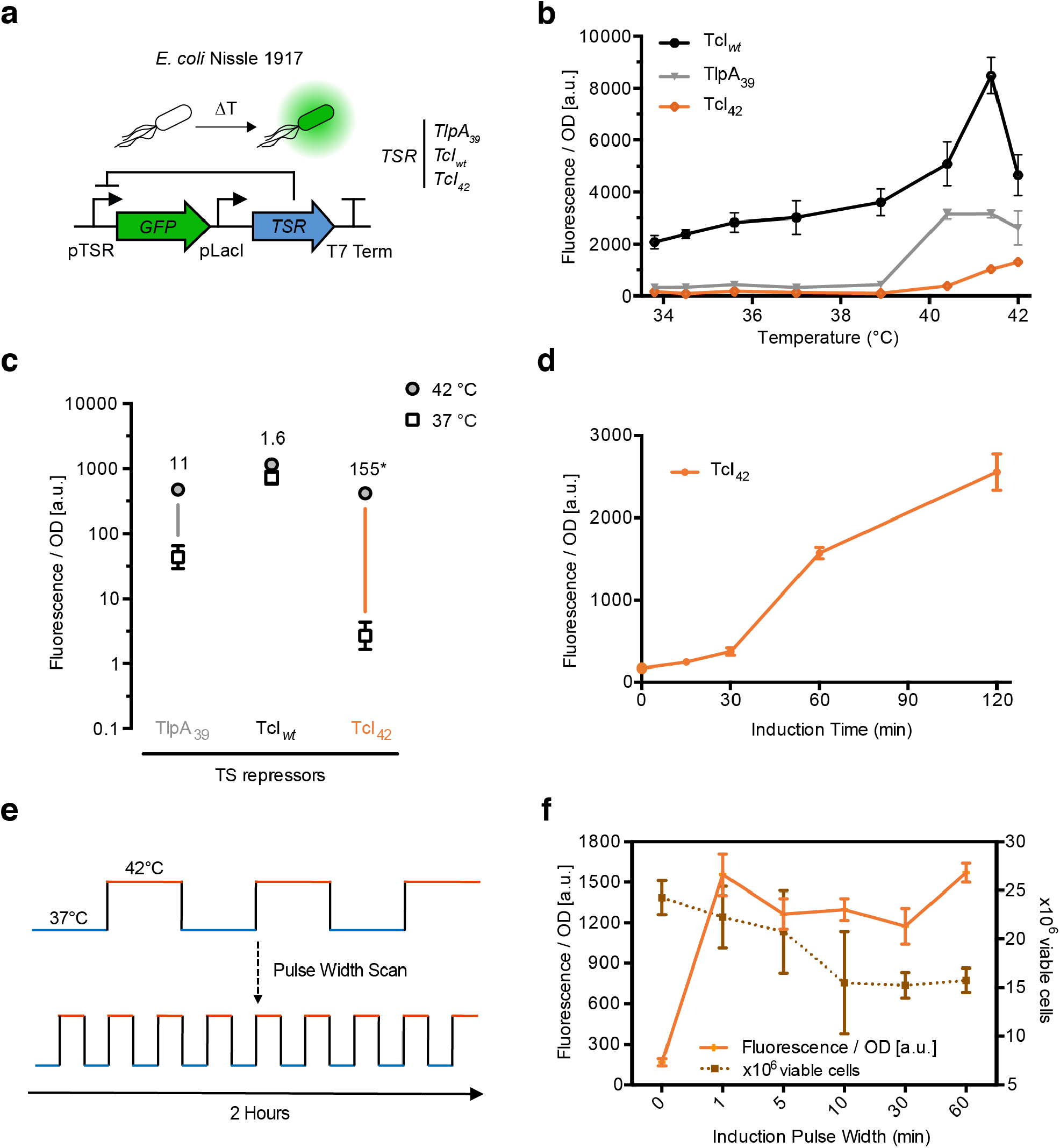
Evaluating temperature-sensitive transcriptional repressors in *E. coli* Nissle 1917. (**a**) Illustration of the genetic circuit used to characterize the behavior of temperature-sensitive repressors in *E. coli* Nissle 1917. (**b**) Optical density (OD_600_)-normalized fluorescence as a function of induction temperature for a fixed duration of 1 hour, measured 24 hours after induction. (**c**) OD-normalized fluorescence 24 hours after a 1-hour induction at 37 °C or 42 °C for the constructs shown in **b**. Numbers indicate fold-change. (**d**) OD-normalized fluorescence as a function of induction duration. Cells were stimulated at 42 °C and fluorescence measured 24 hours later. (**e**) Illustration of the pulsatile heating scheme used to optimize thermal induction and cell viability. (**f**) OD-normalized fluorescence as a function of pulse duration for the TcI_42_ circuit. All samples were stimulated for a total of 1 hour at 42 °C and 1 hour at 37 °C and evaluated 24 hours later. Viable cell counts at various pulse durations plotted to reflect cell viability. Where not seen, error bars (±SEM) are smaller than the symbol. N=4 biological replicates for each sample.

Results from these experiments indicated that TcI42 is the best candidate for integration into our thermal switch since it exhibits strong induction at 42 °C while maintaining low levels of baseline activity.

With TcI42 serving as the thermal transducer in our cells, we next sought to determine the minimal heating duration and ideal heating parameters required to achieve strong activation while minimizing damage to cells. We stimulated cells carrying the circuit described in **Fig. 1a** by elevating the temperature to 42 °C for different durations and measured the corresponding fluorescence intensity (**Fig. 1d**). The results indicated that a minimal heating time of one hour is needed for robust activation. We quantified the effect of this thermal dose on microbial cell viability and simultaneously tested a pulsatile heating scheme that was previously shown to enhance viability in mammalian cells^35^. For the pulsatile heating scheme, the duty cycle was kept constant at 50% while alternating the temperature between 37 °C and 42 °C, resulting in a total of one hour at 42 °C over a two-hour period, with pulse duration varying between 1 and 60 minutes (**Fig. 1e**). As hypothesized, cell viability decreased as the pulse duration increased, while induction levels did not significantly vary (**Fig. 1f**). Based on these results, we selected a five-minute pulse duration for subsequent applications, as this heating paradigm enhanced cell viability while being readily achievable with a focused ultrasound setup. Collectively, our experiments identified and characterized TcI42 as an effective thermal transducer to control gene expression in the therapeutically relevant EcN strain.

### Constructing a thermally actuated state switch

On its own, the TcI42 switch is not sufficient for microbial cancer therapy. This switch is transiently activated for the duration of heating, while tumor therapy requires weeks to effectively suppress tumor growth. Since daily FUS application over this period is infeasible in a clinical setting, we set out to engineer a gene circuit that maintains a prolonged therapeutic response following a single, brief thermal activation.

To enable stable thermal switching, we placed the expression of Bxb1, a serine integrase, under the control of a thermally inducible promoter regulated by TcI42 (**Fig. 2a**). Our design combines the temperature sensitivity of TcI42 with the permanent effector function of the Bxb1 integrase. At physiological temperatures of approximately 37 °C, constitutively expressed TcI42 represses the expression of Bxb1. Upon thermal stimulation, the release of TcI42 repression results in a burst of Bxb1 expression. Thermally derepressed Bxb1 expression catalyzes the inversion and activation of a P7 promoter that is responsible for driving the expression of a fluorescent reporter to monitor the state of the circuit and a tetracycline resistance cassette serving as a placeholder for a therapeutic protein (**Fig. 2a**). Because the DNA inversion event that activates P7 is permanent, this promoter will continue to drive the expression of its protein payloads even when the temperature stimulus is terminated. To avoid unregulated expression of Bxb1 we insulated the activity of the temperature-activated promoter by inserting two strong terminators upstream to block activity from other regions of the plasmid^36^.

**Figure 2.**
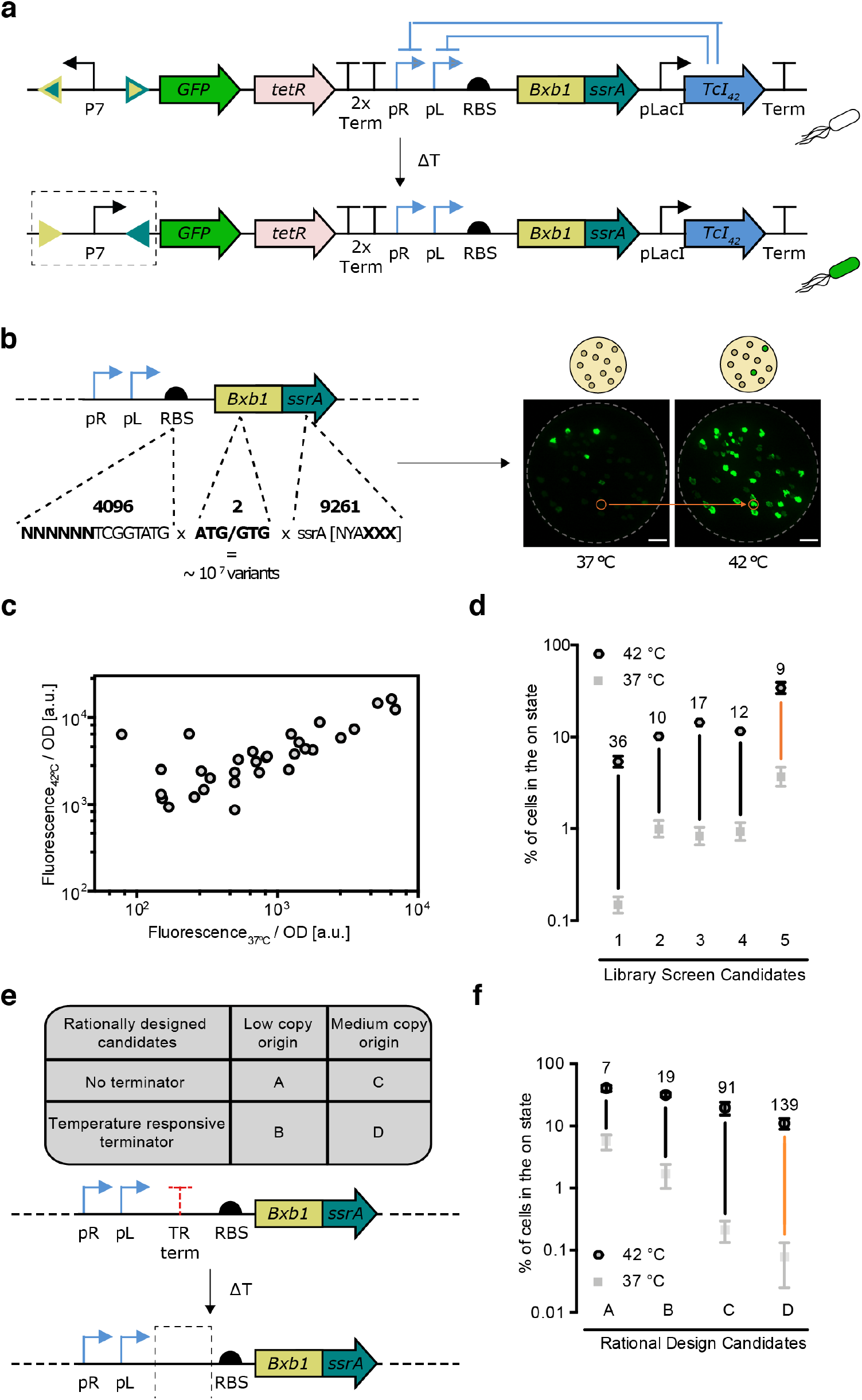
Construction and optimization of a temperature responsive state switch. (**a**) Illustration of the genetic circuit constructed to establish a temperature responsive state switch. TetR is the tetracycline resistance cassette. (**b**) Illustration of the sites targeted in a high throughput screen to optimize circuit switching. A representative fluorescence image of replica plates used to screen for circuit variants. Plates were incubated at the indicated temperature for one hour and further incubated at 37 °C until colonies grew large enough for analysis. The orange circle indicates an example colony selected for further assay. (**c**) Circuit variants from the screen in **b** characterized for their fluorescence at 37 °C and 42 °C. (**d**) Percent conversion to the on-state 24 hours after a 1-hour thermal stimulation at 42 °C or 37 °C for five of the circuit variants from (**c**). (**e**) Summary of rational modifications made to reduce leakage in the circuit at 37 °C. (**f**) Percent induction 24 hours after a 1-hour of thermal induction at 42 °C compared to baseline incubation at 37 °C for four circuit variants described in **e**. Error bars represent ± SEM. N=4 biological replicates for each sample.

The ideal performance of the circuit described above would maintain low baseline activity at physiological temperature while providing strong and lasting induction once thermally stimulated. To achieve this performance, we tuned three key sequence elements affecting Bxb1 translation and stability: the Bxb1 ribosomal binding sequence (RBS), start codon, and ssrA degradation tag (**Fig. 2b**). To efficiently identify the best versions of these elements we performed a library screen that consisted of randomized 6-bp sequences within the Bxb1 RBS, two Bxb1 start codon choices, and randomized terminal tripeptides in the Bxb1 ssrA degradation tag^37^. Two start codons were tested because the non-canonical start codon GUG can down-regulate ribosomal efficiency, and the last three amino acids of the ssrA degradation tag were randomized because they strongly modulate the degradation rate of ssrA-tagged proteins^38^. A total landscape of approximately 10^7^ possible unique variants was sampled using a high-throughput plate-replication assay (**Fig. 2b**). Agar plates containing colonies of library members were first replicated, and then one plate was incubated at 37 °C to assess baseline expression, while the other plate was stimulated at 42 °C for an hour and returned 37 °C for the rest of the growth period. The temperature-dependent fluorescence of a representative sampling of variants is shown in **Figure 2c**. We selected a subset of variants with low leak and high activation to quantify their switching performance with a larger number of replicates (**Fig. 2d**). Out of these candidates, we selected candidate #5 for further optimization since it activated the largest percentage of the cells upon stimulation, a metric that is important to ensure strong therapeutic activity *in vivo*, while still retaining a reasonable temperature-dependent fold change (**Fig. 2d**).

To reduce the baseline activity of candidate #5, we modified two additional circuit components (**Fig. 2e**). The first modification changed the origin of replication from the low-copy origin pSC101 to the medium-copy origin p15A. The second modification explored the effect of inserting a temperature-sensitive terminator upstream of the Bxb1 coding sequence. This terminator introduces a temperature-sensitive secondary structure in the mRNA transcript that helps terminate protein expression at low temperatures, adding to the control provided by TcI42 to prevent leaky Bxb1 protein production at physiological temperature^39^. At 42 °C, this terminator loses its secondary structure and Bxb1 expression is unimpeded. We assessed the performance of four constructs with either one or both of these modifications (**Fig. 2f**). Increasing the copy number of the plasmid and inserting the terminator reduced baseline activation independently. When combined together, these modifications resulted in significantly reduced leakage while maintaining a large fold-change in activated cells upon induction. The resulting construct, obtained through a combination of randomized and rational engineering, displayed a more than 100-fold change in activity between 37 °C and 42 °C.

### Engineering cells for thermally actuated secretion of anticancer immunotherapy

To demonstrate that engineered cells containing our optimized thermally actuated circuit can function in a clinically relevant scenario, we modified the output of the circuit to express a therapeutic payload **(Fig. 3a)**. In particular, we selected αCTLA-4 and αPD-L1 nanobodies, which block signaling through the CTLA-4 and PD-L1 checkpoint receptor pathways, that are heavily implicated in T-cell silencing within immunosuppressive solid tumors. Checkpoint inhibitors such as αCTLA-4 and αPD-L1 have emerged as a major class of cancer therapy, but their therapeutic efficacy is commonly accompanied by the risk of unintentionally activating autoimmunity in bystander tissues when administered systemically^40,41^. By combining the ability of FUS to target specific areas deep within tissues with a highly specific thermal switch, we reasoned that we could target the activity of these potent immunomodulators to tumors and thereby mitigate the risk of systemic exposure.

**Figure 3.**
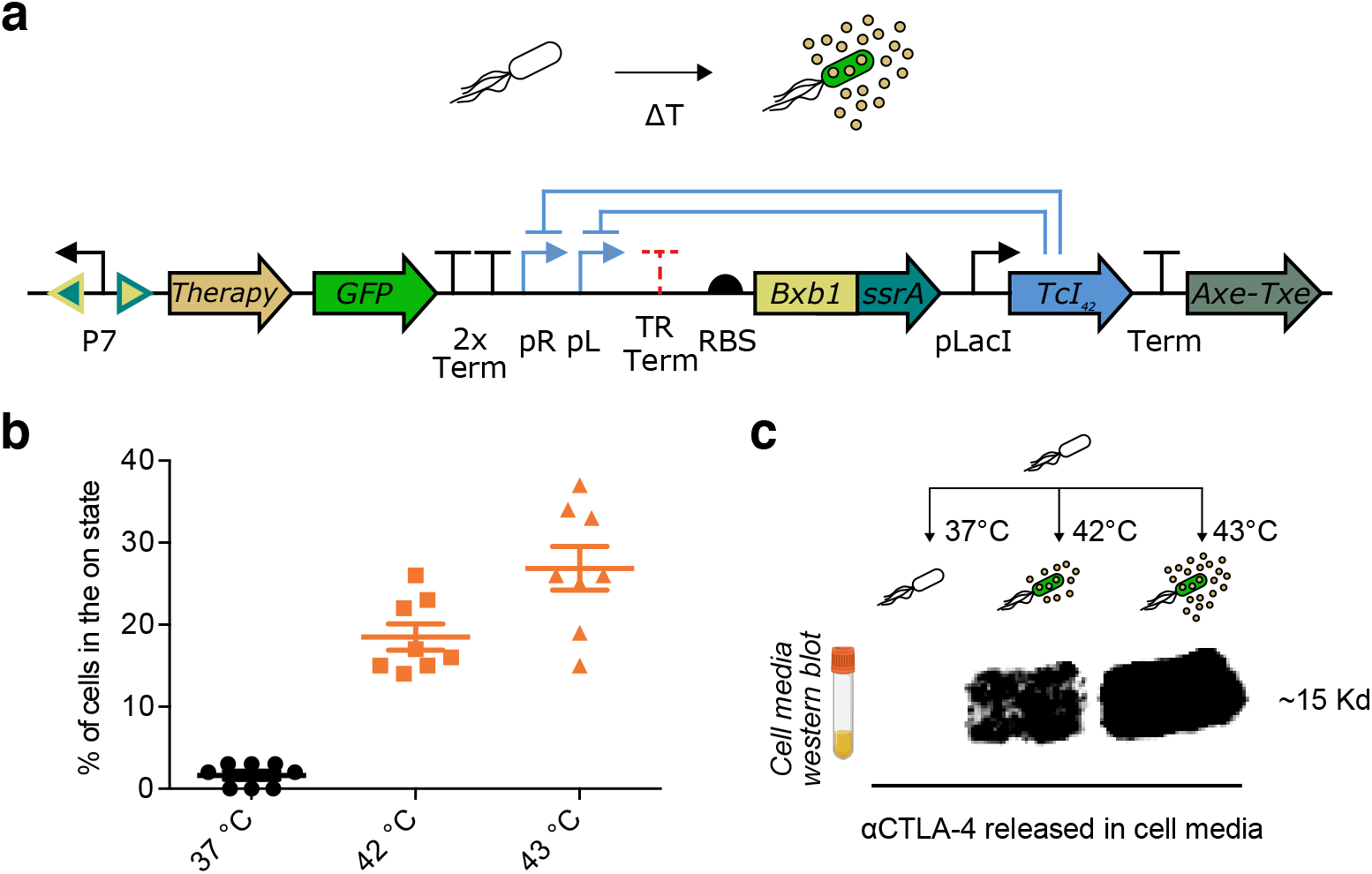
Thermally activated sustained release of immunotherapeutic payload. (**a**) Temperature responsive state switch modified to release αCTLA-4 or αPD-L1 nanobodies. The circuit includes an Axe-Txe stability cassette. (**b**) Percent activation 24 hours after a 1-hour of thermal induction at 37 °C, 42 °C or 43 °C for the circuit described in **a**. (**c**) Western blot against hexahistidine-tagged αCTLA-4 nanobodies. Cells were induced for 1 hour at 37 °C, 42 °C or 43 °C, then expanded in 5 ml of media for 24 hours at 37 °C before collecting the media and assaying for the release of αCTLA-4 nanobodies. The original western blot image is shown in **Fig. S1**. Similar staining was done to confirm αPD-L1 release.

αCTLA-4 and αPD-L1 have been shown to produce antitumor effects when released by tumor-injected probiotics^10^. We hypothesized that local release of these proteins in tumors by FUS-activated systemically administered bacteria would supress tumor growth. To test this hypothesis, we fused αCTLA-4 and αPD-L1 to a PelB secretion tag to enhance their secretion upon activation and cloned each construct in place of the tetracycline cassette in our optimized switching circuit. In addition, to stabilize our plasmids for long-term retention *in vivo* without antibiotic selection, we added an Axe-Txe toxin-antitoxin stability domain, which ensures retention of the plasmid in a cell population by eliminating cells that lose it^42,43^.

The thermal switching functionality of our therapeutic circuits closely resembled their non-therapeutic counterpart. The circuit containing αCTLA-4 maintained a tight off-state at 37 °C while exhibiting robust fold-changes upon induction at 42 °C and 43 °C **(Fig. 3b)**. The 43 °C condition was added to allow for comparison with *in vivo* experiments, where this slightly higher temperature is targeted as the focal maximum inside the tumor to allow more of the mass to be heated above 42 °C.

To assess the secretion of therapeutic nanobodies upon activation, we stimulated the cells for one hour at 37 °C, 42 °C and 43 °C, then cultured them for one day at 37 °C, and performed a Western Blot to evaluate the levels of αCTLA-4 nanobodies released in their media. This experiment demonstrated that αCTLA-4 nanobodies are reliably secreted exclusively upon stimulation at 42 °C and 43 °C **(Fig. 3c)**. We could not detect any secretion when the cells were incubated at 37 °C. Similar characterization was performed for cells expressing αPD-L1.

### Activation of engineered microbes with focused ultrasound elicits in vivo tumor suppression

To enable thermal control of engineered therapeutic microbes *in vivo* we built a FUS stimulation setup capable of locally delivering an activation signal within tumors. An ideal setup should be able to elevate the local temperature within a tumor to a predetermined level and autonomously cycle between that temperature and 37 °C every five minutes to enact our optimized pulsatile heating scheme. In our FUS hyperthermia system **(Fig. 4a)**, a holder secured an anesthetized tumor-bearing mouse vertically in a degassed water chamber. The chamber also held a submerged feedback-controlled ultrasound transducer that used the temperature of the tumor to adjust its output intensity in real time to achieve the target temperature. We demonstrated that this system is capable of toggling the temperature in the tumor of a living animal between 37 °C and 43 °C every five minutes **(Fig. 4a)**. We set the focal maximum temperature inside the tumor at 43 °C to allow more of the mass to be heated above 42 °C and ensure reliable activation within the context of a mouse. While this could lead to some thermal damage, we reasoned that such damage within the tumor is acceptable and could help enhance the microbial therapy^44,45^.

**Figure 4.**
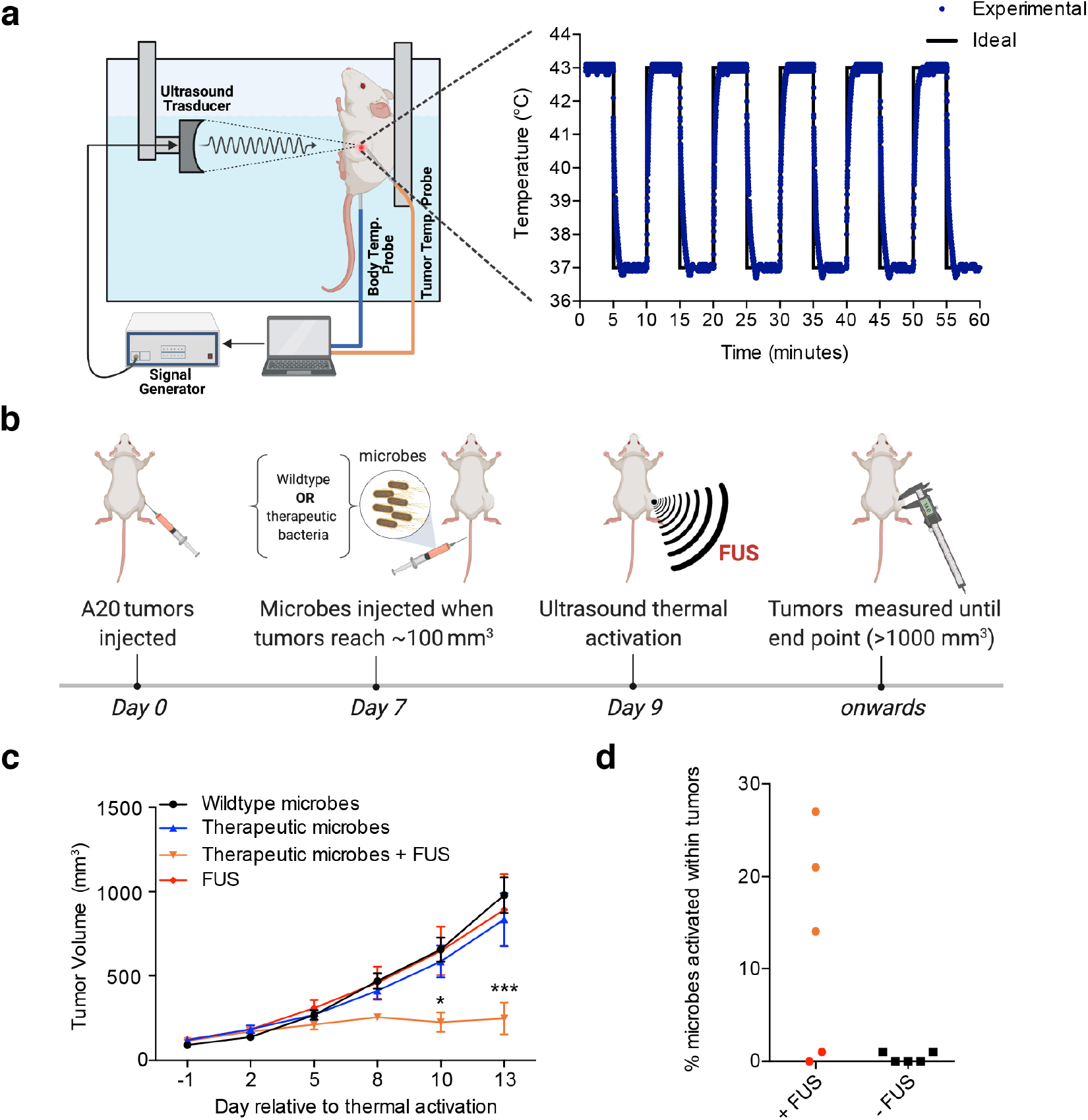
Ultrasound-activated bacterial immunotherapy reduces tumor growth *in vivo*. (**a**) Illustration of the automated setup used to deliver FUS hyperthermia to tumors (left) and representative time course of tumor temperature from a mouse treated with alternating 5-min steps between 37 °C and 43 °C. (**b**) Diagram illustrating the experiment performed to assess the activation of microbial antitumor immunotherapy *in vivo*. Mice were injected with a 1:1 mixture of EcN cells carrying the αCTLA-4 or αPD-L1 circuits, or wildtype EcN. EcN cells were washed and adjusted to 0.625 OD_600_ before injecting 100 μL per mouse intravenously. Ultrasound was applied for a total of 1 hour at 43°C with 50% duty cycle and 5-min pulse duration. (**c**) Tumor sizes measured over two weeks in mice treated with wildtype EcN, therapeutic microbes in the absence of FUS, therapeutic microbes and FUS treatment, or FUS treatment alone. (**d**) Percent activation of therapeutic EcN isolated from FUS-treated and non-FUS-treated tumors two weeks after FUS treatment. One of the FUS-activated tumors disappeared after treatment and bacterial activation inside it could not be quantified. Where not seen, error bars (±SEM) are smaller than the symbol. At least five mice were used for each control condition.

Using this *in vivo* setup, we tested our ability to locally activate systemically administered therapeutic microbes inside tumors. We seeded 5×10^6^ A20 murine tumor cells in the right flanks of BALB/c mice **(Fig. 4b)**. Once the tumors grew to approximately 100 mm^3^, we intravenously injected 10^8^ EcN cells comprising a 1:1 mixture of cells engineered for thermally-actuated αCTLA-4 or αPD-L1 secretion. Injected microbes were given two days to engraft in tumors before they were stimulated with FUS. After FUS activation, tumor growth was monitored to assess therapeutic efficacy.

We observed major retardation in tumor growth in FUS-treated tumors colonized by therapeutic cells, while growth rates in controls including non-FUS treated mice, animals treated with only FUS, and subjects injected with wild-type EcN were substantially higher **(Fig. 4c)**. After completing this experiment, we collected the tumors, chemically homogenized them, and plated the suspension on selective media. By counting the percentage of activated bacteria, we demonstrated that our thermal switch is exclusively activated in targeted tumors and remains active up to at least two weeks post activation **(Fig. 4d)**. One of the six FUS-activated tumors disappeared as a result of the treatment and bacterial activation inside it could not be quantified. In two out of six FUS-treated tumors, ultrasound failed to activate the therapeutic bacterial circuit. This could be due to limitations in our heating setup, which is currently capable of only partially heating the tumor mass. The two non-activated mice were removed from our analysis of tumor growth. Overall, our *in vivo* experiments demonstrated that EcN cells engineered for thermally controlled checkpoint inhibition could home to and engraft in tumors from systemic circulation, become activated specifically in response to FUS, maintain this activity for at least two weeks after a 1-hour FUS treatment and significantly reduce tumor growth.

## DISCUSSION

Our results establish a cell-based system for targeted immunotherapy that couples the special ability of therapeutic bacteria to home into the necrotic core of solid tumors with the capacity of FUS to locally activate their therapeutic function. The sustained activation of these therapeutic bacteria is enabled by a thermal state switch developed through high throughput genetic engineering to have low baseline activity, rapid induction upon stimulation and sustained activity *in situ*. When this state switch is used to actuate the release of immune checkpoint inhibitors, the resulting engineered microbes can be activated inside tumors by brief FUS exposure to secrete their therapeutic payload over an extended timeframe and substantially reduce tumor growth.

The growing body of work on bacteria-based therapies^46–49^ and the increasing clinical acceptance of FUS^50,51^ provide FUS-actuated bacterial therapeutics a path to ultimate clinical implementation. Potential disease targets include cancers with readily identified primary masses that are challenging to resect surgically, such as head-and-neck, ovarian, pancreatic or brain tumors. FUS-actuated bacterial therapeutics could be also relevant to metastatic tumors since microbial therapy in a single tumor mass can generate a strong systemic adaptive immune response, and eliminate distant tumor lesions through a potent abscopal effect^52^. However, further work will be needed to optimize the timing, dose and molecular identity of FUS-activated therapy release for each application. To enhance therapeutic efficacy, it may be beneficial to combine FUS-activated bacterial therapeutics with other molecular or cellular therapies. For example, engineered bacteria and immune cells have distinct and often complementary tumor entry and engraftment profiles. Engineering microbes that successfully enter immunosuppressed tumor regions to secrete checkpoint inhibitors or cytokines could help make this environment more accessible to engineered T cells. In this way, the bacteria and T cells can synergistically exert their therapeutic function from the inside-out and from the outside-in, respectively. Beyond tumor therapy, locally activated bacterial agents have potential utility in a wide array of other biomedical applications. For example, FUS-controlled state switches could be useful in controlling the activity of gut microbes *in vivo*^53^, the function of cell-based living materials *in vitro*^54–57^, and in industrial metabolic engineering^24,58^.

## MATERIALS AND METHODS

### Plasmid Construction and Molecular Biology

All plasmids were designed using SnapGene (GSL Biotech) and assembled via reagents from New England Biolabs for KLD mutagenesis (E0554S) or Gibson Assembly (E2621L). After assembly, constructs were transformed into NEB Turbo (C2984I) and NEB Stable (C3040I) *E. coli* for growth and plasmid preparation. The Bxb1 recombinase-encoding gene was a kind gift of Richard Murray (Caltech). Integrated DNA Technologies synthesized other genes and all PCR primers. Plasmids containing the αCTLA-4, αPD-L1, and Axe-Txe genes were kind gifts from of Tal Danino (Columbia University).

### Preparation of cell lines for in vitro and in vivo experiments

Plasmids containing engineered genetic circuits were transformed into Nissle 1917 *E. coli* (Mutaflor®). Nissle cells were cultured in LB broth (Sigma) and grown on LB agar plates (Sigma) containing appropriate antibiotics. Singular colonies were picked into LB broth and grown overnight in a shaking incubator (30 °C, 250 rpm). The next day, optical density measurements (OD_600_) were taken, and the saturated cultures were diluted to 0.1 OD_600_. Diluted cultures were then allowed to grow to exponential phase until they reached 0.6 OD_600_ before starting assays. Optical density measurements were taken using a Nanodrop 2000c (Thermo Scientific) in cuvette mode.

### Western Blot

Five milliliters of cell media were collected for each sample and concentrated with an Amicon® Ultra-15 Centrifugal Filter Unit. Concentrated cell media was then mixed with Laemmli loading buffer and BME before loading into a pre-cast polyacrylamide gels SDS-PAGE gel (Bio Rad) and ran at 75 V for 140 minutes. Western blotting was performed using the Transblot Turbo apparatus and nitrocellulose membrane kit (Bio Rad). Transfer was performed at 25 V for 7 minutes. Membranes were blocked with 5% Blotto milk (Santa Cruz Biotechnology) in 0.05% TBS-Tween for 1 hour at room temperature. Primary staining was performed using the mouse anti-His sc-8036 antibody (Santa Cruz Biotech) overnight at 4 °C. Blots were then washed three times for 15 minutes at 4 °C with 0.05% TBS-Tween and stained for 4 hours with mouse IgG kappa binding protein (m-IgGκ BP) conjugated to Horseradish Peroxidase (HRP) (Santa Cruz Biotech, sc-516102) at room temperature. After three 15-minute washes, HRP visualization was performed using Super signal west Pico PLUS reagent (Thermo Fisher Scientific). Imaging was performed in a Bio-Rad ChemiDoc MP gel imager. A subsequent epi white light image of the blot under the same magnification was acquired to visualize the stained molecular weight standards.

### Thermal regulation assay

Once bacterial cell cultures reached approximately 0.6 OD_600_, 50 μL aliquots of each sample was transferred into individual Bio-Rad PCR strips with optically transparent caps and subsequently heated in conditions specific to the experiment using a Bio-Rad C100 Touch thermocycler with the lid set to 50 °C. Following heating, cells continued to incubate overnight undisturbed at either 30 °C (Figure 1) or 37 °C (Figure 2-4). The PCR strips were then removed, vortexed, and spun down, and the green fluorescence of each of the samples was measured using the Strategene MX3005p qPCR (Agilent) and an unamplified FAM filter. To measure cell density, the samples were diluted 1:4 with fresh LB media (without antibiotic) and then transferred into individual wells of a 96-well plate (Costar black/clear bottom). Optical density measurements were taken using the SpectraMax M5 plate reader (Molecular Devices). In order to quantify the temperature-dependent gene expression (*E*) using background-subtracted, OD-normalized fluorescence (Fig. 1b-d, 1f, 2c), Equation (1) was used:

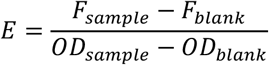

In this equation, we define *F* as the raw fluorescence measurement and *OD* is the OD_600_ measurement of the sample. The value of the blank fluorescence and blank optical density was determined as the average of *N = 4* samples of untransformed Nissle cells, as opposed to engineered Nissle cells, in LB.

### Screens to optimize circuit behavior

To improve Bxb1 thermal regulation, a sequence randomized library of the RBS, start codon, and ssrA degradation tag was ordered from Integrated DNA Technologies. PCR products that included the Bxb1 coding region and immediately surrounding sequences were amplified using custom primers and were inserted into the backbone of the rest of the parent plasmid using Gibson Assembly (Fig. 2b-d). This library was transformed into EcN and plated on LB Agar plates with antibiotic resistance at a low colony density of approximately 30 colonies per petri dish. Following overnight incubation at 30 °C to allow the colonies to become visible, these plates were then replicated into two daughter petri dishes using a replica-plating tool (VWR 25395-380). The parent petri dish was incubated at 4 °C until the conclusion of the experiment. One daughter plate was grown overnight at the baseline temperature of 37 °C, and the other was incubated at 42 °C for 1 hour and then moved to 37 °C overnight. After colonies became visible, the plates were imaged using a 530/28 nm emission filter to determine colonies that were fluorescent at the ‘on’ temperature but opaque at the ‘off’ temperature (Bio-Rad ChemiDoc MP imager). Promising library variants were then picked from the corresponding parent petri dish at 4 °C and analysed against the parent plasmid of the library using the liquid culture fluorescence-based assay described above.

### Percent switching assay

Strips of liquid bacteria samples were prepared and incubated in the Bio-Rad Touch thermocycler. After the prescribed thermal stimulus and incubation at 37 °C, PCR strips were removed, vortexed, and spun down on a tabletop centrifuge. Five 1:10 serial dilutions in liquid LB were then performed, transferring 10 μL of sample into 90 μL of LB media sequentially. After thorough mixing, 50 μL of the most diluted samples was plated onto an LB plate and allowed to incubate at 30 °C overnight. Upon the appearance of visible colonies, plates were imaged using the same Bio-Rad ChemiDoc MP imager with both blue epifluorescence illumination and the 530/28 nm emission filters. The percentage of colonies in the ‘on state’ (*P*) was determined according to Equation (2):

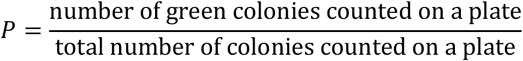

### Animal procedures

All animal procedures were performed under a protocol approved by the California Institute of Technology Institutional Animal Care and Use Committee (IACUC). 8-12 week-old BALB/c female mice were purchased from Jackson Laboratory. To establish A20 tumor models in mice, 5 × 10^6^ A20 cells were collected and suspended in 100 μL phosphate buffer saline (PBS) prior to subcutaneous injection into the flank of each mouse. When tumor volumes reached approximately 100 mm^3^, engineered EcN cells prepared according to the procedure outlined in the section above were then collected by centrifugation (3000 g for 5 min), washed with phosphate buffer saline PBS 3 times, and diluted in PBS to 0.625 OD_600_. 100 μL of the resulting solution was injected into each of the A20 tumor bearing mice via tail vein. For thermal actuation using ultrasound, mice were anesthetized using a 2% isoflurane-air mixture and placed on a dedicated animal holder. Anesthesia was maintained over the course of the ultrasound procedure using 1-1.5% isoflurane, adjusted in real-time to maintain the respiration rate at 20-30 breaths per minute. Body temperature was continuously monitored using a fiber optic rectal thermometer (Neoptix). When appropriate, the target flank was thermally activated using the automated FUS setup described below, cycling between the temperatures of 43 °C and 37 °C every 5 minutes for 1 hour of total heating. Following ultrasound treatment, the mouse was returned to its cage and the size of its tumor was measured with a caliper to track the therapeutic efficacy. When the tumors reached ∼1000 mm^3^ mice were culled and the tumors were collected for analysis. Mice that did not have microbial cells in their tumors were excluded from the study.

### Tumor analysis

Tumors were collected and homogenized in ten milliliters of PBS containing 2 mg/ml collagenase and 0.1 mg/ml DNAse for one hour at 37 °C. Homogenized tumors were serially diluted and plated onto LB plates to quantify the number of cells colonizing the tumors. The percentage of cells activated within tumors was determined by counting the number of GFP positive cells.

### Feedback-controlled focused ultrasound

We developed a closed loop thermal control setup to maintain a specified predetermined temperature within the tumor of a mouse by modulating the intensity of the FUS. This setup includes a water bath filled with pure distilled water that is being actively cleaned and degassed with an AQUAS-10 water conditioner (ONDA) and maintained at 33 °C with a sous vide immersion cooker (InstantPot Accu Slim). A tumor-bearing mouse that has been anesthetized as described above is fastened nose up vertically to an acrylic arm that is connected to a manual 3D positioning system (Thorlabs) to enable 3D motion of the mouse within the water bath. A Velmex BiSlide motorized positioning system is used to submerge and position the 0.67 MHz FUS transducer (Precision Acoustics PA717) such that the focal point of the transducer lies within the tumor of the mouse. A signal generator (B&K #4054B) generates the thermal ultrasound signal which is then amplified (AR #100A250B) and sent to drive the ultrasound transducer. The water in this chamber acts as the coupling medium to transfer the ultrasound wave from the transducer to the tumor. To measure the internal tumor temperature during a heating session we temporarily implant a thin fiber optic temperature probe (Neoptix) into the tumors. This temperature readout is also used to align the focus of the transducer with the tumor by emitting a constant test thermal ultrasound signal. Once the system is aligned, we run a Matlab closed loop thermal control script that regulates the signal generator output. Feedback for the controller is provided by the temperature measurements acquired with a sampling rate of 4 Hz. The actuator for the controller is the voltage amplitude of the continuous sinusoidal signal at 0.67 MHz used to drive the FUS transducer, where the voltage is adjusted also at 4 Hz. The system uses a PID controller with anti-windup control that modifies the amplitude of the thermal ultrasound waveform to achieve a desired temperature in the targeted tissues. The Kp, Ki, Kd, and Kt parameters for the PID and anti-windup were tuned using Ziegler-Nichols method, and in some cases adjusted further through trial-and-error tuning to achieve effective thermal control.

### Statistics and replicates

Data is plotted and reported in the text as the mean ± S.E.M. Sample size is *N = 4* biological replicates in all *in vitro* experiments unless otherwise stated. This sample size was chosen based on preliminary experiments indicating that it would be sufficient to detect significant differences in mean values. *P* values were calculated using a two-tailed unpaired *t*-test.

### Data and code availability

Plasmids will be made available through Addgene upon publication. All other materials and data are available from the corresponding author upon reasonable request.

## ACKNOWLEDGEMENTS

The authors thank Tal Danino, Tiffany Chien, Candice Gurbatri, Sreyan Chowdhury, Dan Piraner and Victoria Hsiao for sharing reagents and helpful discussions. Figure 4b was created with BioRender.com. This research was funded by the Sontag Foundation, the Army Institute for Collaborative Biotechnologies (W911NF-19-D-0001) and the Defence Advanced Research Projects Agency (D14AP00050). M.H.A. was supported by the NSF graduate research fellowship and the Paul and Daisy Soros Fellowship for New Americans. AB-Z was supported by the European Union’s Horizon 2020 research and innovation programme under the Marie Skłodowska-Curie grant agreement No. 792866. Related research in the Shapiro laboratory is supported by the Burroughs Welcome Career Award at the Scientific Interface, the Packard Foundation Fellowship in Science and Engineering, the Pew Scholarship in the Biomedical Sciences and the Heritage Medical Research Institute.

## AUTHOR CONTRIBUTIONS

M.H.A. and M.G.S. conceived the study. M.H.A., M.S.Y, D.R.M, M.S and AL-G planned and performed experiments. D.R.M. wrote the MATLAB script for *in vivo* thermal control. AB-Z and D.R.M helped with building the ultrasound heating setup. M.H.A. and M.S.Y analysed data. M.H.A., M.S.Y, D.R.M and M.G.S. wrote the manuscript with input from all other authors. M.G.S. supervised the research.

