## Supplemental Figure 1 for "Acoustic Remote Control of Bacterial Immunotherapy"

**SUPPLEMENTARY FIGURES**

**a**

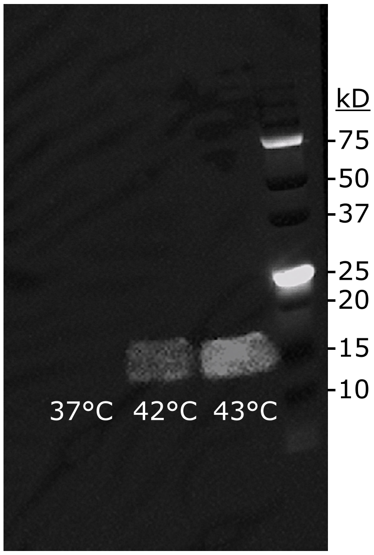


**Figure S1 | Western blot to assay for the release of αCTLA-4 upon thermal activation.** (**a**) Unmodified image of the western blot shown in **Fig. 3c.** The image in Fig. 3c was cropped and inverted to make it fit better into the figure presentation.
